# Vascular epiphytes contribute disproportionately to global centres of plant diversity

**DOI:** 10.1101/2021.05.21.445115

**Authors:** Amanda Taylor, Gerhard Zotz, Patrick Weigelt, Lirong Cai, Dirk Nikolaus Karger, Christian König, Holger Kreft

**Affiliations:** Biodiversity, Macroecology & Biogeography, Faculty for Forest Sciences & Forest Ecology, University of Göttingen, Büsgenweg 1, 37077 Göttingen, Germany; Institute of Biology and Environmental Sciences, Carl von Ossietzky University of Oldenburg, Ammerländer Heerstrasse 114, 26129 Oldenburg, Germany; Smithsonian Tropical Research Institute, Apartado Postal 0843-03092, Balboa, Ancon, Panamá, República de Panamá; Swiss Federal Research Institute WSL, Zürcherstrasse 111, 8903 Birmensdorf, Switzerland; Ecology and Macroecology group, Institute for Biochemistry and Biology, University Potsdam, Potsdam, Germany; Centre of Biodiversity and Sustainable Land Use (CBL), University of Goettingen, Büsgenweg 1, 37077 Göttingen, Germany

**Keywords:** epiphytes, latitudinal diversity gradient, Neotropics, Orchidaceae, pteridophytes, tropical forest

## Abstract

**Aim:** Vascular epiphytes are ubiquitous features of wet tropical forests where they contribute substantially to local and regional plant diversity. While some basic epiphyte distribution patterns are relatively well studied, little effort has been made to understand the drivers responsible for constraining their global distribution. This study quantifies the substantial contribution of epiphytes to global gradients and centres of vascular plant diversity and explores whether epiphytes vary from terrestrial plants in relation to contemporary and historical environmental variables.

**Location:** Global.

**Time period:** Present.

**Major taxa studied:** Vascular epiphytes.

**Methods:** We integrated EpiList 1.0, a comprehensive list comprising > 30,000 vascular epiphyte species, and species distributions derived from the GIFT database to describe the global biogeography of epiphytes. We used generalized linear mixed effects models to assess the relationship between epiphytic and terrestrial plant diversity, and contemporary and historical environmental predictors.

**Results:** We find that epiphytes substantially contribute to global centres of vascular plant diversity, accounting for up to 39% of the vascular flora in the Neotropics. Epiphytes decrease in species numbers with increasing latitude at a rate three times faster than terrestrial plants, a trend that is driven mainly by the distribution of tropical forests and precipitation. Further, large regional differences emerge that are explained by several large endemic angiosperm families (e.g., Neotropical Bromeliaceae) that are absent in other tropical regions.

**Main conclusions:** Our results show that epiphytes are disproportionately diverse in most global centres of plant diversity and play an important role in driving the global latitudinal diversity gradient for plants. The distribution of precipitation and tropical forests emerge as major drivers of the latitudinal diversity gradient in epiphyte species richness. Finally, our findings demonstrate how epiphyte floras in different biogeographical realms are composed of different families and higher taxa revealing an important signature of historical biogeography.

## Introduction

Vascular epiphytes – defined as non-parasitic plants that germinate and are permanently structurally dependent on other plants – are one of the most prominent life forms in tropical forest canopies (Zotz, 2013b). In humid tropical forests, epiphytes may locally account for up to 50% of the vascular flora (Kelly *et al.*, 2004), while globally they constitute roughly 10% of the world’s plant biodiversity (Zotz *et al.*, 2021). Where they reach higher abundances, epiphytes play a critical role in forest nutrient and water cycling (Gotsch *et al.*, 2016), and can contribute substantially to local plant biomass (Díaz *et al.*, 2010; Nadkarni *et al.*, 2004; Zotz, 2016). Moreover, epiphytes provide crucial habitat for canopy-dwelling fauna (Stuntz *et al.*, 2002; Méndez‐Castro *et al.*, 2018), while also adding to the structural complexity of forest canopies (Zotz, 2016).

Globally, some basic patterns in epiphyte richness are well known: epiphytes reach their greatest numbers in the tropics and decrease in numbers towards the poles (Madison, 1977; Gentry & Dodson, 1987; Benzing, 1990; Zotz, 2016), epiphyte richness is lower in the north-relative to the southern hemisphere (Dawson, 1986; Gentry & Dodson, 1987; Zotz, 2005; Burns, 2010), and distribution patterns of epiphytic pteridophytes differ from those of epiphytic seed plants (Madison, 1977; Zotz, 2016). Despite a relatively good understanding of epiphyte distribution patterns, no study has explored such patterns at a global scale for over 30 years (Gentry & Dodson, 1987), and little progress has been made to understand the mechanisms responsible for constraining the global distribution of epiphytes. Since the last global assessment, the advancement of molecular phylogenetic techniques and the development of key spatial databases has considerably improved both plant taxonomic classification and our knowledge of plant distributions, including epiphytes (Zotz *et al.*, 2021). Now it is possible to fully grasp the global extent of vascular epiphyte distributions, addressing questions on how epiphytes contribute to overall vascular plant diversity or whether their responses to environmental gradients differ compared to terrestrial plants.

Recent studies have illustrated striking differences in epiphyte diversity patterns compared to terrestrial representatives from certain groups (e.g., pteridophytes, Nervo *et al.*, 2016; orchids, Taylor *et al.*, 2021), which suggests that epiphytic and terrestrial plants indeed vary in their responses to environmental gradients. Epiphytes are expected to be more tightly coupled to atmospheric conditions than terrestrial plants due to their aerial growth habit and limited access to water, which strongly influences the vertical partitioning of epiphytes within the canopy (Mendieta-Leiva *et al.*, 2020). Evidence for water availability being typically the most important limiting factor for epiphytes is provided by the frequent occurrence of traits among different epiphytic lineages related to water capture, storage, and use-efficiency, such as water-impounding tanks (Zotz *et al.*, 2020), fleshy leaves, succulent stems (including pseudobulbs, Göbel *et al.*, 2020), crassulacean acid metabolism (CAM) photosynthesis (Benzing, 1987), or specialised aerial roots that aid to capture and store water (Pridgeon, 1987).

Epiphytism has evolved independently multiple times among vascular plants, and the evolution of epiphytism-associated traits is thought to have prompted the rapid and independent diversification observed in some plant families, such as the Orchidaceae (75% epiphytic, Zotz *et al.*, 2021; Silvera *et al.*, 2009; Givnish *et al.*, 2015) Bromeliaceae (59% epiphytic, Zotz *et al.*, 2021; Givnish *et al.*, 2011), and leptosporangiate ferns (e.g., Polypodiaceae 89% epiphytic, Hymenophyllaceae 72% epiphytic, Zotz *et al.*, 2021; Schuettpelz & Pryer, 2009). Still, epiphytism is highly unevenly distributed throughout the plant kingdom, being prevalent in some clades, and surprisingly under-represented (e.g., Asteraceae, Poaceae, <0.1% epiphytic, gymnosperms <0.2%, Zotz *et al.*, 2021) or absent from others (e.g., Brassicaceae, Euphorbiaceae, Fabaceae). The uneven representation of epiphytes among clades also emerges geographically. For instance, while there is a relatively uniform representation of epiphytic families across all tropical realms, there is a considerable disparity in the number of species (Gentry & Dodson, 1987). Why epiphytism evolved in a similar number of plant families in different regions, yet only diversified in some regions is not fully understood, although numerous hypotheses have been proposed (Table 1).

**Table 1.**
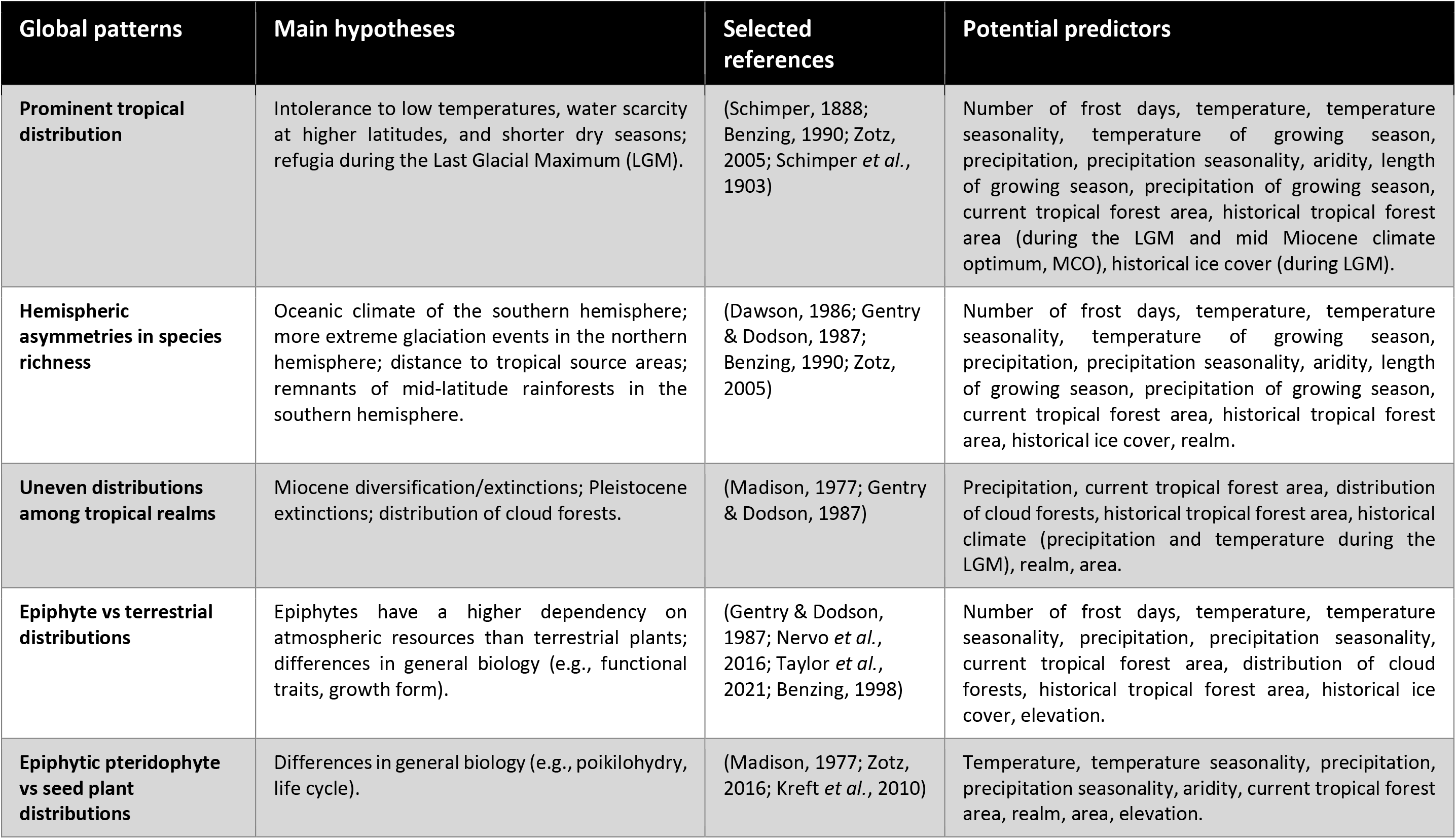
Summary of observed global patterns in epiphyte richness and associated hypotheses, references, and potential predictors.

| Global patterns | Main hypotheses | Selected references | Potential predictors |
| --- | --- | --- | --- |
| **Prominent tropical distribution** | Intolerance to low temperatures, water scarcity at higher latitudes, and shorter dry seasons; refugia during the Last Glacial Maximum (LGM). | (Schimper, 1888; Benzing, 1990; Zotz, 2005; Schimper *et al.*, 1903) | Number of frost days, temperature, temperature seasonality, temperature of growing season, precipitation, precipitation seasonality, aridity, length of growing season, precipitation of growing season, current tropical forest area, historical tropical forest area (during the LGM and mid Miocene climate optimum, MCO), historical ice cover (during LGM). |
| **Hemispheric asymmetries in species richness** | Oceanic climate of the southern hemisphere; more extreme glaciation events in the northern hemisphere; distance to tropical source areas; remnants of mid-latitude rainforests in the southern hemisphere. | (Dawson, 1986; Gentry & Dodson, 1987; Benzing, 1990; Zotz, 2005) | Number of frost days, temperature, temperature seasonality, temperature of growing season, precipitation, precipitation seasonality, aridity, length of growing season, precipitation of growing season, current tropical forest area, historical tropical forest area, historical ice cover, realm. |
| **Uneven distributions among tropical realms** | Miocene diversification/extinctions; Pleistocene extinctions; distribution of cloud forests. | (Madison, 1977; Gentry & Dodson, 1987) | Precipitation, current tropical forest area, distribution of cloud forests, historical tropical forest area, historical climate (precipitation and temperature during the LGM), realm, area. |
| **Epiphyte vs terrestrial distributions** | Epiphytes have a higher dependency on atmospheric resources than terrestrial plants; differences in general biology (e.g., functional traits, growth form). | (Gentry & Dodson, 1987; Nervo *et al.*, 2016; Taylor *et al.*, 2021; Benzing, 1998) | Number of frost days, temperature, temperature seasonality, precipitation, precipitation seasonality, current tropical forest area, distribution of cloud forests, historical tropical forest area, historical ice cover, elevation. |
| **Epiphytic pteridophyte vs seed plant distributions** | Differences in general biology (e.g., poikilohdry, life cycle). | (Madison, 1977; Zotz, 2016; Kreft *et al.*, 2010) | Temperature, temperature seasonality, precipitation, precipitation seasonality, aridity, current tropical forest area, realm, area, elevation. |

One possible explanation for the poorer representation of epiphytes outside of the tropics is that most epiphytic lineages evolved under warm, humid, and non-seasonal conditions (Benzing, 1989) and the evolution of novel functional strategies to withstand a desiccating environment have remained relatively constant through time (i.e., *niche conservatism*, Wiens *et al.*, 2010). Thus, functional traits to persist in very cold or highly seasonal climates (e.g., geophytic habit, annual life cycle) are generally not compatible with the epiphytic life form, making it difficult for individuals to survive outside of tropical habitats. Given that most epiphytic lineages evolved in tropical forests, and since past conditions leave a legacy in contemporary diversity patterns (Sandel *et al.*, 2020), the cooler, drier conditions during the Pleistocene and past distributions of tropical forest biomes may have also played a role in driving global patterns of epiphyte richness, although contemporary distributions of tropical forests are also important (Dawson & Snedon, 1969; Madison, 1977; Gentry & Dodson, 1987, Table 1).

Here, we present the most detailed and quantitative assessment of the global distribution of epiphytes to date, which aims at disentangling spatial patterns of epiphyte richness and their contribution to global centres of plant diversity. We integrate a comprehensive epiphyte list (EpiList 1.0, Zotz *et al.*, 2021) with distribution information for 27,850 epiphyte species obtained from numerous literature sources (Table S3) as well as the Global Inventory of Floras and Traits database (Weigelt *et al.*, 2020). First, we present a global map of epiphyte species richness, highlighting the relative contribution of different families to continental and global patterns of epiphyte diversity. Second, we re-evaluate prominent hypotheses that have been proposed to explain epiphyte distribution patterns (Table 1), including both historical e.g., past distributions of forest biomes, glaciation events, past climatic conditions (Gentry & Dodson, 1987; Zotz, 2005; Dawson, 1986), and contemporary drivers e.g., current distribution of forest biomes, current climatic conditions, elevational range (Kreft *et al.*, 2004; Krömer *et al.*, 2005). Lastly, we compare patterns of epiphyte richness to terrestrial plant richness and establish whether they differ in their responses to environmental conditions.

## Methods

### Epiphyte and terrestrial plant distribution data

As a baseline list of all known epiphyte species, we used the EpiList 1.0 database, which contains over 31,000 epiphyte and hemiepiphyte species names collated from 978 literature sources (Zotz *et al.*, 2021). We included in our analyses all obligate epiphytes and hemiepiphytes, defining hemiepiphytes as plants that germinate in tree crowns (as epiphytes) but unlike true epiphytes grow roots that eventually make contact with the forest floor (Zotz, 2013a). We justify including hemiepiphytes as they begin life as epiphytes and are thus under the same constraints during the most vulnerable life stage. Species names were standardised according to the World Flora Online taxonomic backbone (WFO, 2019) for seed plants, and the World Ferns database (Hassler, 2021) for ferns and lycophytes (hereafter pteridophytes). To gain complete, global coverage of epiphyte distributions, we derived distribution data from a variety of literature and database sources. All pteridophyte species distributions were obtained from the World Ferns database (Hassler, 2021), while seed plant distributions were mainly derived from the Global Inventory of Floras and Traits database (GIFT, Weigelt *et al.*, 2020), and the World Checklist of Selected Plant Families (WCSP, 2018). External literature searches were performed for species that could not be matched to any database.

To obtain a continuous global scheme of regional epiphyte composition, we aggregated the number of epiphyte species following the TDWG (Taxonomic Database Working Group) scheme of botanical countries. The TDWG scheme defines botanical countries as standardised geographical boundaries independent of political configurations (Brummit, 2001). Some TDWG regions do not have complete species lists (e.g., ‘Chile South’), although complete lists are available for the smaller, nested regions within. Thus, in cases where complete checklists were available for all smaller regions nested inside a larger TDWG region, we aggregated these (e.g., Chile South = Región de los Lagos, Aisén del General Carlos Ibáñez del Campo, Región de Magallanes y de la Antártica Chilena). While we aimed to have complete global coverage of all epiphyte species distributions, some TDWG regions may only be partially covered due to incomplete checklists, or because checklists only represented certain plant families or nested regions. Despite this, we have complete global coverage for most of the largest epiphyte families, such as Araceae, Bromeliaceae, Orchidaceae, (WCSP, 2018), and most pteridophytes (Hassler, 2021). Our final dataset amounted to a total of 76,427 species distribution records for 27,850 epiphyte species in 276 regions (including continental island and mainland regions). Although oceanic islands can have diverse epiphyte floras (3,608 species from this study), we excluded them because isolation and island ontogeny significantly influence plant assembly and richness patterns (Whittaker *et al.*, 2008).

To compare epiphyte and terrestrial plant distributions, we subtracted the total number of epiphytes per botanical country from the total number of vascular plants to obtain the total number of terrestrial species per botanical country and the proportional representation of epiphytes (hereafter ‘epiphyte quotient’, sensu Hosokawa, 1950). The same procedure was completed for two subsets - pteridophytes and seed plants. This required selecting botanical countries with complete pteridophyte, seed plant, and total richness values, which was not possible for every botanical country. Thus, our subset for the epiphyte quotient analysis was smaller, including a total of 267 regions for pteridophytes, 195 for seed plants, and 196 for the total number of epiphytes.

We corrected for differences in area size among regions by standardising species richness estimates per region to 10,000 km^−2^ following Kier *et al.*, (2005). We did this by first deriving empirically the slope (*z* value) of the global epiphyte species-area relationship assuming the well supported power law model, which was then incorporated into a modified species-area equation:

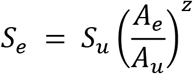

where *S_e_* is the area-corrected species richness of a region, *S_u_* is the regions observed species richness, *A_e_* is the constant area of 10,000 km^2^, A_*u*_ is the observed area of a region in km^2^, and *z* is the slope of the species-area relationship in log-log space (for *z* values see Table S1). We accounted for geographical variation in species-area relationships by estimating separate slopes for each biome (Gerstner *et al.*, 2014). Consequently, Greenland and Aruba were omitted due to being the sole regions in the ‘rock and ice’ and ‘mangrove’ biome, respectively. Biomes were assigned to regions based on the scheme defined by Dinerstein *et al.*, 2017, Table S2). Area corrections were only applied for mapping and comparisons of epiphyte richness among regions, with uncorrected area size being used in all models.

### Environmental predictor variables

We related epiphyte richness to 10 contemporary and past environmental variables out of 19 variables that were initially considered. Variables included three contemporary climatic variables derived from CHELSA V1.2 (Karger *et al.*, 2017) – mean annual precipitation (mm, hereafter precipitation), precipitation seasonality (coefficient of variation in precipitation), and mean minimum monthly temperature (°C, hereafter temperature), all of which have been previously hypothesised or shown at regional to global scales to be important predictors of both epiphytic and terrestrial plant species richness (Zotz, 2005; Gentry & Dodson, 1987; Kreft & Jetz, 2007; Taylor *et al.*, 2021). As a measure of habitat availability for epiphytes, we included the contemporary extent of tropical forest biomes (km^2^). Contemporary tropical forest biomes were extracted from a global map of terrestrial biomes (Olson *et al.*, 2001), and overlaid with our botanical country polygons in order to quantify the total area of tropical forest for each botanical country. In addition, we selected area (km^2^, Weigelt *et al.*, 2020) and elevational range (m, from the Global Multi-resolution Terrain Elevation data set by Danielson & Gesch, 2011 at a resolution of 30 arc-seconds), which are important geographical predictors of species diversity.

We further considered three historical factors, reflecting past climate – Last Glacial Maximum (LGM) ice cover (km^2^, Ehlers *et al.*, 2011), and past distribution of tropical forests– LGM tropical forest area (km^2^, Ray & Adams, 2001) and tropical forest area during the Mid-Miocene climate optimum (km^2^, Henrot *et al.*, 2010). The extent of both historical tropical forest biomes was quantified in the same manner as for the distribution of contemporary tropical forested biomes. However, because biome definitions differed between datasets, we first standardised all biomes to match across datasets, delineating “tropical rainforest”, “sub-tropical forest”, and “tropical seasonal forest” to “tropical forest”. Finally, we explore continental differences in epiphyte occurrences, which allow inferences about idiosyncratic historical biogeographic patterns not captured by the environmental predictors (variable biogeographic realm). Similar to the classification of biomes, each region was assigned its respective realm following Olson *et al.*, (2001), in order to explore the richness and relative contribution of different epiphyte families among the different continents. All data used for this analysis can be found in Table S2.

We also considered the number of frost days, length of growing season, precipitation of growing season, mean temperature of growing season, temperature seasonality, aridity, distribution of cloud forests, LGM precipitation, and LGM temperature. These variables were not included in the final analyses due to being highly correlated with other variables (at the threshold of *r* ≥ 0.70), and for showing weaker relationships (lower correlation *r* values) with epiphyte species richness compared to the retained uncorrelated variables (Dormann *et al.*, 2013). For example, the number of frost days was highly correlated with temperature (*r* = 0.86), and we retained temperature due to its stronger relationship with epiphyte species richness (*r* = 0.61) compared to the number of frost days and epiphyte richness (*r* = −0.41).

### Statistical analyses

Generalised linear mixed effects models (GLMMs) with a poisson and binomial error distribution (for proportion data) were implemented to analyse the effects of different environmental drivers on epiphyte and terrestrial plant richness and proportional representation, respectively. We chose GLMM models to overcome overdispersion, which we accounted for by including an observational-level random effect. We fitted separate GLMMs for seed plants and pteridophytes. All predictor variables except for temperature were log10(x+1) transformed to reduce skewness. Moreover, to make the results comparable across different models, we scaled all predictor variables to zero mean and unit variance. Minimum adequate models were selected based on the corrected Akaike Information Criterion (AICc). We considered models with a ΔAICc value < 2.0 compared to the minimum AICc value to be the best supporting models following Burnham and Anderson (2004). Model residuals showed a low degree of spatial autocorrelation (e.g., total epiphyte richness: Moran’s *I* = 0.08, p ≤ 0.01). Including a spatial residual autocovariate (RAC) further reduced (Dormann *et al.*, 2007) spatial autocorrelation (e.g. total epiphyte richness: Moran’s *I* = 0.03, p = 0.06). However, because we found that accounting for spatial autocorrelation did not qualitatively alter our results, we opted to present the non-spatial models.

Quasi-poisson and binomial GLMs were used to assess the simple relationship between epiphyte and terrestrial richness, proportional representation, and absolute latitude. For all proportion models, we did not consider regions with fewer than 3 plant species to reduce distortion of global patterns. Lastly, we regressed our epiphyte richness model residuals with absolute latitude to confirm that we captured all possible combinations of factors explaining variation in epiphyte richness with increasing latitude. All statistical analyses were conducted in the R environment (version 4.0.0, R Core Team, 2020) using the packages jtools (Long, 2019), lme4 (Bates *et al.*, 2015), MuMIn (Barton, 2020), and spdep (Bivand & Wong, 2018).

## Results

### The contribution of epiphytes to global vascular plant diversity

Barring Mediterranean climates, epiphytes substantially contribute to global centres of vascular plant diversity, accounting for 39% of the flora of Ecuador (excluding Galapagos), and 23-26% of the floras of Panama, Brazil South, Costa Rica, New Guinea, and Colombia, respectively. Similarly, we find marked differences in epiphyte richness patterns, with area-corrected epiphyte richness being most impressive in the Neotropics, particularly in Ecuador (1699 species per 10,000 km^-2^; 5574 unstandardized total species richness), Costa Rica (1454; 2660), Panama (1192; 2513), and Colombia (958; 5520). On the contrary, epiphytes are under-represented in regions with large deserts (e.g., Egypt) or frequent freezing temperatures (e.g., Central European Russia, Figure 1).

**Figure 1.**
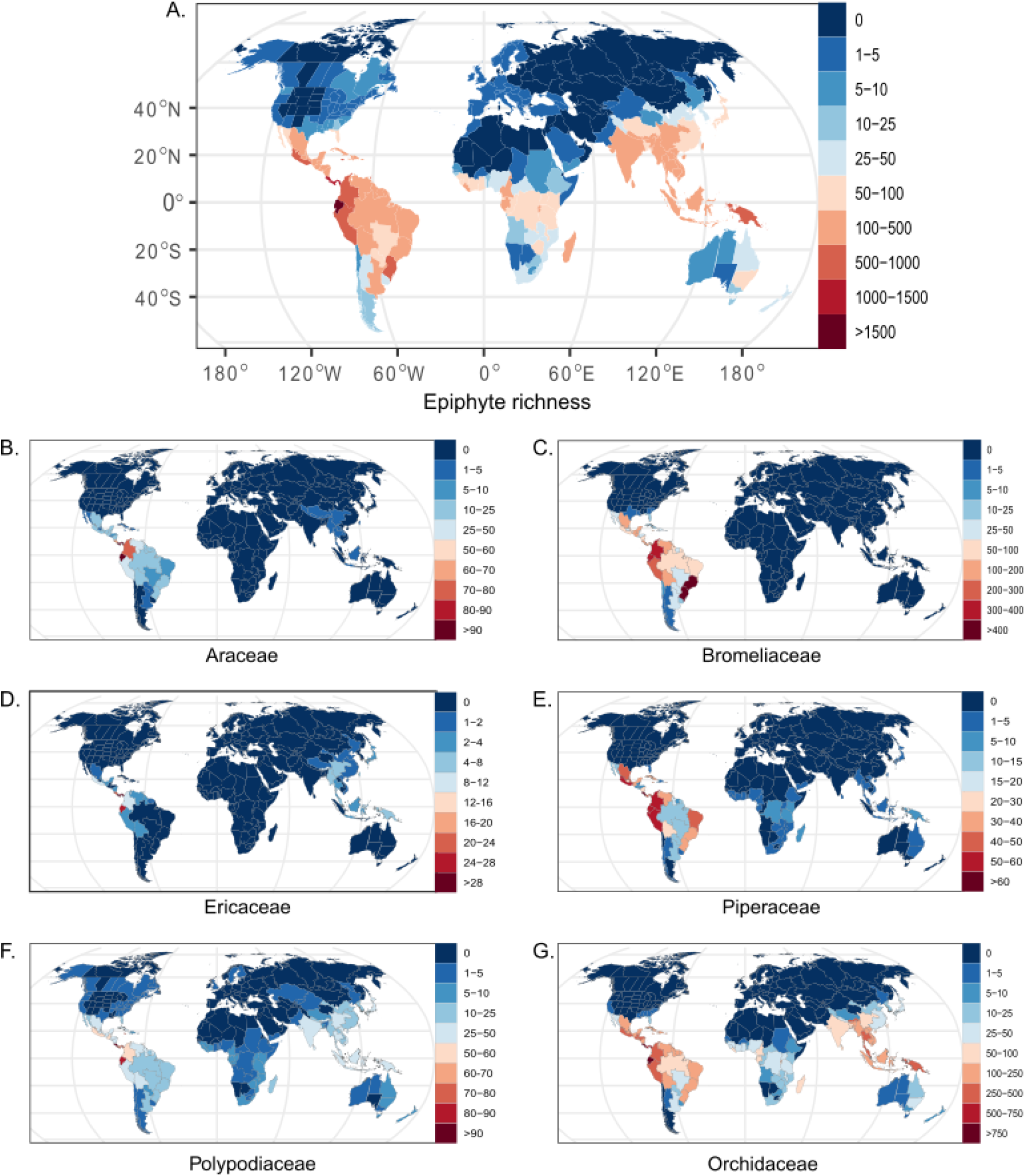
Global patterns of area-corrected epiphyte species richness per 10,000 km^−2^, (A) for all vascular epiphytes and for six of the most species-rich epiphyte families; (B) Araceae, (C) Bromeliaceae, (D) Ericaceae, (E) Piperaceae, (F) Polypodiaceae, and (G) Orchidaceae.

As well as contributing to global centres of vascular plant diversity, epiphytes also play an important role in driving the latitudinal diversity gradient (Figure 2). Specifically, epiphytes decrease in species numbers with increasing absolute latitude at a rate 3× faster than terrestrial plants (epi slope: −1.50 ± 0.14; terr slope −0.42 ± 0.05; p = ≤ 0.01, Figure S1). Further highlighting the role of epiphytes in driving the latitudinal diversity gradient, we find that while the epiphyte quotient (% epiphytes) decreases with increasing latitude, the proportion of terrestrial species increases (% epi slope: −1.33 ± 0.14; % terr slope: 1.33 ± 0.01; p = ≤ 0.01, Figure 2A). We also note a pronounced latitudinal asymmetry in which epiphyte richness and quotient decreases from the tropics to higher latitudes almost twice as more rapidly in the northern-relative to the southern hemisphere. This relationship is stronger for epiphytic seed plants (north slope: −1.54 ± 0.01; south slope: −0.91 ± 0.01; p = ≤ 0.01) than for pteridophytes (north slope: −0.92 ± 0.02; south slope: −0.17 ± 0.02; p = ≤ 0.01), whose distributions extend further into temperate regions (Figure 2B,C).

**Figure 2.**
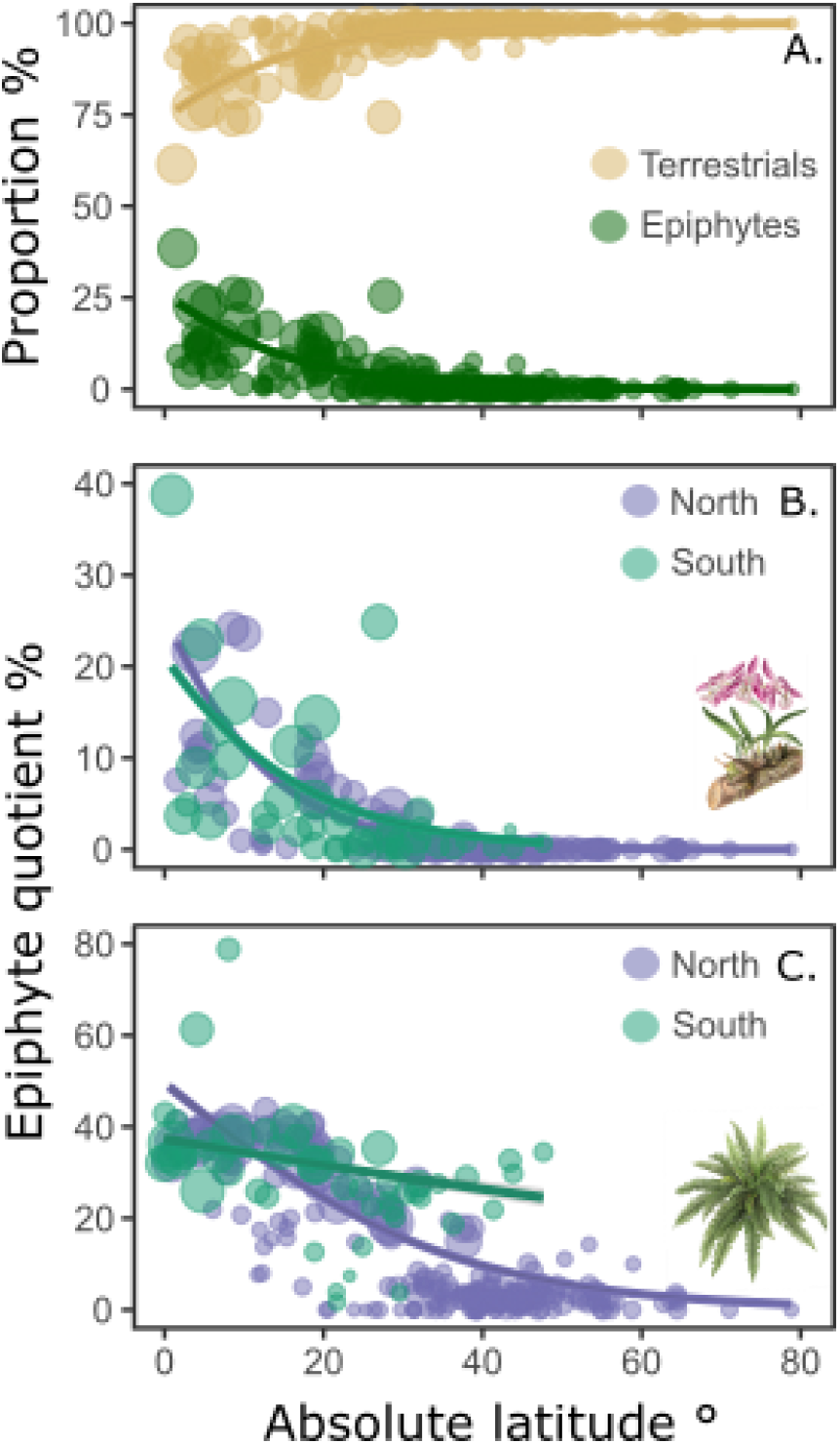
Latitudinal gradients in (A) the proportion of epiphyte (green) and terrestrial (gold) plant species, and latitudinal asymmetries for epiphyte quotients % of (B) seed plants and (C) pteridophytes between the northern- (purple) and southern hemispheres (green). Points indicate regions weighted by species richness, with larger points indicating higher species richness. Lines indicate the strength of the relationship, including 95% confidence intervals.

Unsurprisingly, the Neotropics emerge as the most diverse biogeographical realm, with 51 out of 68 epiphytic plant families represented (Figure 3), including 63% of all epiphytes in our study (17,433 epiphyte species, uncorrected for area), and 3.5 times more species than the second most diverse realm (Indomalayan – 4,984 species; 18% of all epiphytes). We find this pattern to be robust even when controlling for differences in area, climate and other variables among regions (Figure 4). Conversely, the Afrotropics is the least diverse tropical realm (1,714 species; 6%), containing fewer species than Australasia (4,602; 17%), with a notable absence or poor representation of several large angiosperm families, including Bromeliaceae (0 species), Ericaceae (0 species), Araceae (1 species), and Gesneriaceae (6 species), which form important components of the epiphyte flora in other tropical realms. The two largely temperate Palearctic and Nearctic have the least diverse epiphyte floras, totally 1,219 (4%) and 786 (3%) species, respectively. This pattern also holds when controlling for environmental factors, with the Palearctic and Nearctic having significantly fewer epiphytes, particularly among seed plants, compared to all other biogeographical realms.

**Figure 3.**
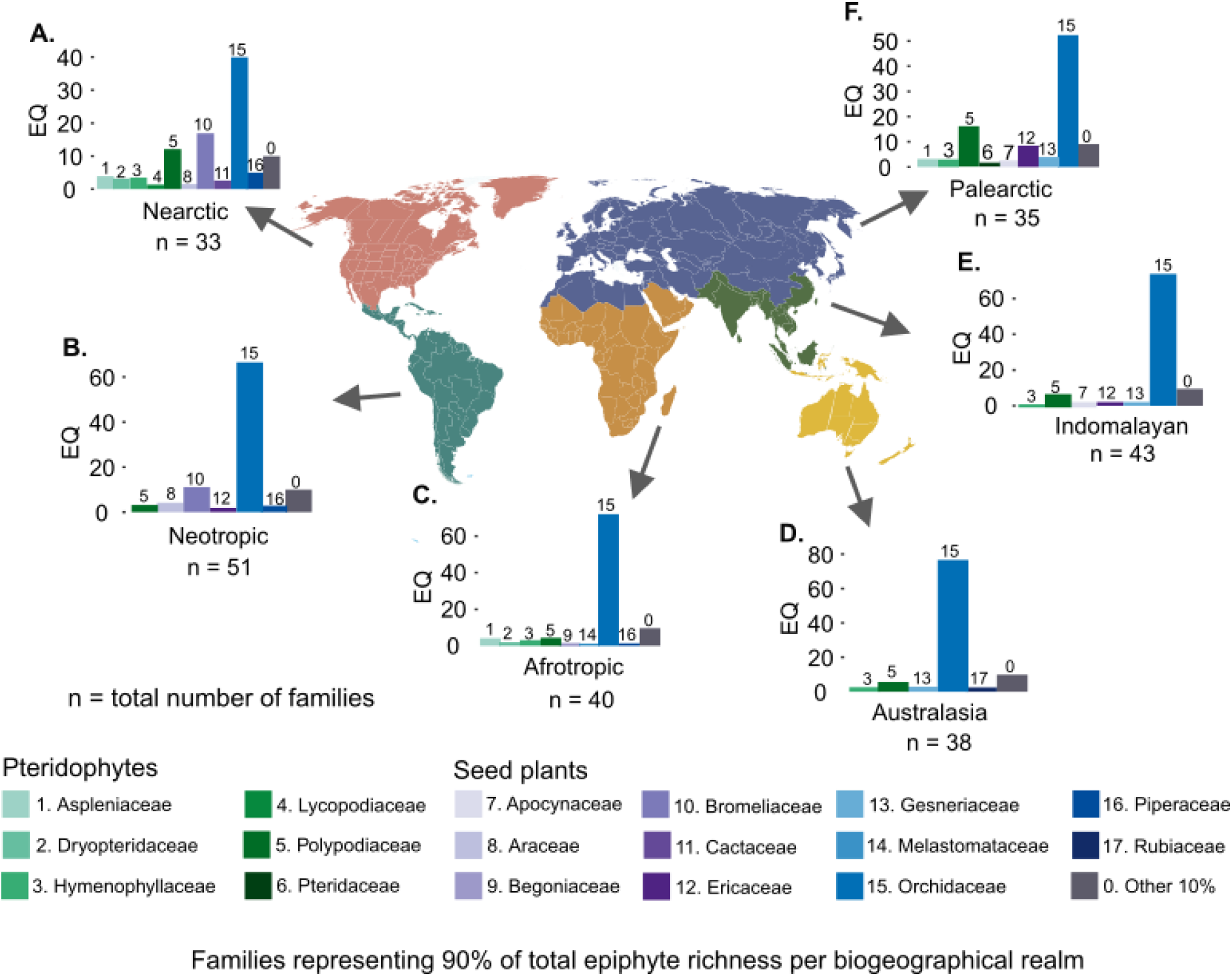
Epiphyte quotient % (EQ) of major families (representing 90% of total epiphyte richness) among different biogeographical realms. Biogeographical realms are defined following Olson *et al.*, (2001). Numbers above each column correspond to families (e.g. 15 = Orchidaceae). Numbers below the names of biogeographical realms indicates the number of families present in that realm.

**Figure 4.**
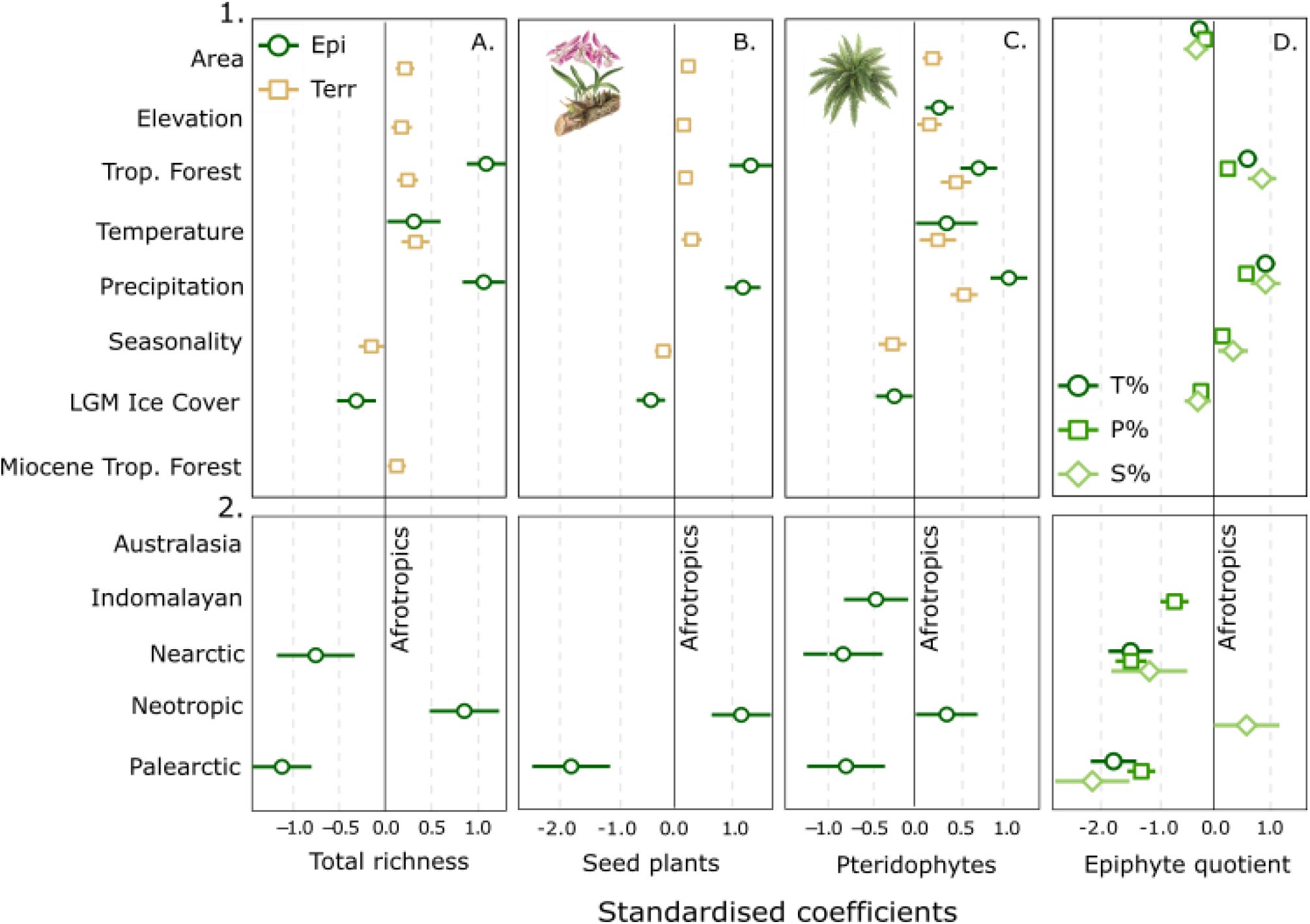
Standardised coefficient plots showing the effect of region area (Area, km2), elevational range (Elevation, m), tropical forest area (Trop. Forest, km^2^), mean minimum monthly temperature (Temperature, °C), mean annual precipitation (Precipitation, mm), precipitation seasonality (Seasonality, CV of Precipitation), ice cover during the last glacial maximum (LGM Ice Cover, km^2^), and tropical forest area during the Mid-Miocene Thermal Optimum (Miocene Trop. Forest, km^2^) on the total number of (A) epiphyte and terrestrial species, and separately for (B) seed plants, and (C) pteridophytes. Only significant predictors are shown. Epiphyte coefficients are indicated with dark green circles and terrestrial plants with gold squares. Panel (D) illustrates the effect of the same set of predictors on the epiphyte quotient % of the total number of epiphytes (T %, dark green circles), pteridophyte epiphytes (P %, green squares) and seed plant epiphytes (S %, light open diamonds). Confidence intervals (95%) are also shown. The predicted values of epiphyte richness and quotient % per biogeographical realm can be found in row 2, where “Afrotropics” is the reference realm (i.e., 0 = Afrotropics).

Low epiphyte richness does not imply a low epiphyte quotient. To the contrary, the Afrotropics with the lowest epiphyte richness has some of the highest proportions of epiphytic pteridophytes, particularly in Gabon, Equatorial Guinea, and Rwanda, where 41-43% of all pteridophytes are recorded as growing epiphytically. As to be expected, Orchidaceae are the most diverse plant family in all biogeographical realms, ranging in proportional representation from 40% (Nearctic) to 77% (Australasia) with Polypodiaceae being a distant second (3-16%). However, we also observe differences in the contribution of epiphyte families among the biogeographical realms. For example, almost entirely endemic to the New World, Bromeliaceae contribute significantly to epiphyte diversity in the Neotropics (11%) and Nearctic (17%), second only to the Orchidaceae. Likewise, Ericaceae form an important component of the Palearctic epiphyte flora (8%) and Apocynaceae and Gesneriaceae are diverse in the Indomalayan and Australaisian realms (each 3%), respectively.

### Epiphytes and terrestrial plants respond differently to past and present environmental variables

Epiphytes have, on average, higher absolute standardized regression coefficients (slopes) than terrestrial plants, indicating stronger associations with environmental gradients (Figure 4). We find strong positive relationships between epiphyte richness and quotient %, and tropical forest area (epi rich: 1.02 ± 0.10; quotient %: 0.73 ± 0.10) and precipitation (epi rich: 1.10 ± 0.10; quotient %: 0.89 ± 0.10), and a weaker, positive association with epiphyte richness and temperature (epi rich: 0.28 ± 0.13). Ice cover during the Last Glacial Maximum (LGM) also proves to be an important driver of epiphyte richness, revealing a negative relationship (epi rich: −0.26 ± 0.09, Figure 4A). In comparison to epiphytes, terrestrial plants showed stronger positive relationships with geographical variables, such as area and elevation, the latter of which was also positively associated with epiphyte pteridophyte richness (Figure 4C). We also note a surprising result of precipitation seasonality, which is very weakly negatively associated with terrestrial but not epiphytic plant species richness. The combination of environmental predictors, including biogeographical realm, explained a high degree of variance in both the global epiphyte (92%) and terrestrial richness models (79%), and to a lesser extent the total epiphyte quotient model (87%).

Regardless of taxonomic group, all significant environmental variables have greater effects on epiphyte richness than on terrestrial richness, particularly with respect to tropical forest area for seed plants (epi slope 1.39 ± 0.15; terr slope 0.22 ± 0.05, *p* = ≤ 0.01) and precipitation for pteridophytes (epi slope 1.01 ± 0.11; terr slope 0.48 ± 0.06, *p* = ≤ 0.01). Despite this, our combined predictors explain a high degree of variance in both the epiphyte and terrestrial models for seed plants (variance explained epiphytes: 92%; terrestrial plants: 78%) and pteridophytes (epiphytes: 86%; terrestrial plants: 78%). When considering the epiphyte quotient of different taxonomic groups separately, tropical forest area has the greatest positive effect on seed plants (% slope 1.04 ± 0.15, *p* = ≤ 0.01), and precipitation on pteridophytes (% slope 0.89 ± 0.16, *p* = ≤ 0.01). The combined predictors explain a high degree of variance in both the seed epiphyte quotient model (87%) and pteridophyte quotient model (66%). Finally, during inspection of the global epiphyte model residuals, we found no pattern related to absolute latitude (Figure S2), suggesting that the strong latitudinal decrease in epiphyte richness can be well explained by the environmental predictors included.

## Discussion

Our results reveal that epiphytes are not only important contributors to local species diversity, but are also disproportionately diverse in all global centres of plant diversity, barring Mediterranean biomes (see Barthlott et al., 2007 for comparison). We also find compelling evidence that epiphytes play a major role in driving the global latitudinal diversity gradient for plants, revealing a three times faster decrease in species richness with increasing absolute latitude in comparison to terrestrial species.

Tropical forest area and precipitation emerge as key drivers of the latitudinal gradient in epiphyte diversity, illustrating the importance of tropical forests as habitat for epiphytes. Tropical montane cloud forests are particularly important ecosystems for epiphytes due to their high levels of atmospheric humidity (as clouds or fog) and mild temperatures, allowing for increased water interception and, therefore, a reduction in drought stress (Gotsch *et al.*, 2016; Karger *et al.*, 2021). Although we do not fully capture the distribution of tropical montane cloud forest in this coarse-grained analyses, we can draw comparisons with studies along elevational gradients, which often attribute the high diversity of epiphytes at mid-elevations to the presence and conditions of tropical montane cloud forests (Ding *et al.*, 2016; Acebey *et al.*, 2017). Indeed, regions in our study with expansive cloud forests (e.g., Ecuador, Colombia, New Guinea), also have the highest numbers of epiphytes, while regions that have fewer mountainous regions (e.g., Africa) have depauperate epiphyte floras.

Most epiphyte lineages are thought to have evolved under the dark, humid conditions of ancient tropical forests (Schimper, 1888; Schneider *et al.*, 2004) and possess morphological and physiological traits that reflect this (e.g., relating to capturing and storing water). These functional strategies, together with the rapid decrease in epiphyte diversity with decreasing temperatures, precipitation, tropical forest area, and increasing levels of historical ice cover during the Last Glacial Maximum (LGM) suggests that most epiphytes have not deviated from their ancestral niche (niche conservatism, Wiens *et al.*, 2010), and have not developed physiological tolerances to withstand cooler, drier conditions. The pronounced hemispherical asymmetries in epiphyte diversity, where epiphytes decrease in species numbers more rapidly in the northern-relative to the southern hemisphere further support the niche conservatism hypothesis (Zotz, 2005; Madison, 1977; Gentry & Dodson, 1987; Dawson, 1986). South-temperate forests vary considerably from their northern counterparts, being mainly composed of evergreen species and growing in comparatively mild, oceanic climates (Markgraf *et al.*, 1995; McGlone *et al.*, 2016). These temperate rainforests (e.g., conifer-broadleaf forest in New Zealand, Valdivian temperate forest in Chile, Argentina), are often likened to having a similar multi-tiered structure as tropical forests (Dawson & Snedon, 1969; McGlone *et al.*, 2016), and may explain the higher diversity of epiphytes in southern Australia, New Zealand, Argentina, and Chile, compared to equivalent latitudes in the north.

In favour of the “odd man out” argument (Couvreur, 2015), we find the Afrotropics to be the least diverse tropical realm in terms of epiphytes, with a notable absence of epiphytic taxa in many large, cosmopolitan angiosperm families. For example, we recorded no epiphytic species from the Ericaceae family, one epiphytic Araceae, and only a fraction of orchids compared to the Neotropics and Indomalayan realms. This paucity can be explained by drier conditions during the LGM, with a particularly strong reduction of the area of tropical forest biomes in tropical Africa, leading to reduced habitat for epiphytes, widespread extinctions (Carlucci *et al.*, 2017), and reduced speciation compared to the tropical forests of South America and Asia (Kissling *et al.*, 2012). In comparison, the Neotropical realm has over three times the number of epiphytic species of the second most diverse Indomalayan realm, with Orchidaceae (comprising 67% of the epiphytic flora), Bromeliaceae (11%), and Araceae (4%) being the dominant plant families. Outside of the Neotropics, however, the contribution of families to the epiphytic flora of each biogeographical realm is more heterogeneous. For example, after the three most diverse families Orchidaceae (cosmopolitan), Polypodiaceae (cosmopolitan), and Bromeliaceae (Americas), Ericaceae is the next most prominent family in the Palearctic (8%), Piperaceae in the Nearctic (5%), Aspleniaceae in the Afrotropics (4%), and Apocynaceae and Gesneriaceae in the Indomalayan and Australasian realms (each 3%), respectively. Thus, our study also demonstrates how epiphyte floras in different biogeographical realms are composed of different families and higher taxa revealing an important signature of historical biogeography.

Supporting the hypothesis that epiphytes are more strongly coupled to atmospheric conditions, we show that epiphytes generally display stronger relationships with climatic variables than terrestrial plants. One possible reason for this is the niche partitioning of epiphytes to within-tree microclimatic gradients, by which epiphytes display a remarkable functional variety among different taxonomic groups (Benzing, 2004). For instance, at least 14% of epiphytic pteridophytes (Hymenophyllaceae, Zotz *et al.*, 2021) are poikilohydric and due to their inability to control water-loss are generally confined to humid forests (Proctor, 2012), or the lower trunks of host trees (Zotz & Büche, 2000, but see Krömer & Kessler, 2006). This might explain the stronger, positive effect of precipitation on epiphytic compared to terrestrial pteridophytes given that water, which is a scarce resource in tree canopies, is a requirement for pteridophyte reproduction (Proctor, 2007). Similarly, the stronger effect of precipitation, minimum temperature and tropical forest area on epiphytic seed plants might reflect the differences in traits associated with an epiphytic compared to terrestrial life form. For example, most terrestrial orchids can be classified as geophytes, and therefore have traits (e.g., tubers) that aid survival in highly seasonal environments (e.g., Mediterranean or temperate climates, Taylor *et al.*, 2021), while epiphytic orchids have traits more associated with capturing and storing water within the canopy. Thus, terrestrial plants are more heavily constrained by soil conditions, which may confound the signal of macroclimate variables to some degree. Indeed, the coarse-grained nature of our analyses inevitably leads to an under-estimation of the importance of regional variation such as over elevational gradients.

In summary, our study quantifies the substantial role of epiphytes - a generally neglected group of plants - to global centres of plant diversity, highlighting an important role of tropical forest biomes and historical biogeography. However, many questions still remain. Why, for example, is epiphytism so unevenly distributed among plant families? And how does this interact with historical biogeography in determining modern richness gradients? What are the key vegetative and reproductive traits that promote epiphytism in epiphyte-rich families? How does diversification in epiphytic lineages vary in space and time? Our study makes a step in this direction by providing a first quantitative baseline and by illustrating that epiphytes show remarkable differences in diversity patterns compared to terrestrial plants on a global scale and across different taxonomic groups.

## Author contributions

AT, GZ, HK, PW conceptualised the study, GZ assembled the epiphyte database, HK, PW and CK assembled the GIFT database, AT assembled missing species distributions, AT and PW cleaned data, LC provided tropical forest biome data, DK provided additional climate data, AT performed all analyses and drafted the manuscript. All authors contributed substantially to revisions.

## Funding

AT received support from the German Research Foundation (DFG project number 447332176). HK acknowledges funding by the German Research Foundation in the framework of the DynaCom project. CK acknowledges support from the German Science Foundation (DFG, grant no. ZU 361‐1/1). DK acknowledges funding from the Swiss Federal Research Institute for Forest, Snow and Landscape Research internal grant exCHELSA, ClimEx, the Joint BiodivERsA COFUND Call on ‘Biodiversity and Climate Change’ (project ‘FeedBaCks’) with the national funder Swiss National Foundation (20BD21_193907), the ERA-NET BiodivERsA– Belmont Forum with the national funder Swiss National Foundation (20BD21_184131) (part of the 2018 Joint call BiodivERsA–Belmont Forum call; project ‘FutureWeb’), and the Swiss Data Science Projects SPEEDMIND and COMECO.

## Data accessibility statement

We confirm that with the acceptance of the manuscript, all supporting data will be submitted as a supplementary.csv file.

## Supplementary Material

**Table S1.** Slopes (z value) of species-area relationships in log-log space for each biome used to account for geographical variation when calculating area-corrected species richness for the total number of epiphytes (Total), epiphytic pteridophytes (Pter.), seed plants (Seed), Araceae (Arac.), Bromeliaceae (Brom.), Ericaceae (Eric.), Orchidaceae (Orch.), Piperaceae (Pipe.), Polypodiaceae (Poly.).

| Biome | z<br>Total | z<br>Pter. | z<br>Seed | z<br>Arac. | z<br>Brom. | z<br>Eric. | z<br>Orch. | z<br>Pipe. | z<br>Poly. |
| --- | --- | --- | --- | --- | --- | --- | --- | --- | --- |
| Boreal Forests/Taiga | -0.22 | -0.23 | 0.02 | 0 | 0 | 0 | 0 | 0 | -0.10 |
| Deserts & Xeric Shrublands | 0.23 | 0.22 | 0.10 | 0.03 | 0.02 | 0.02 | 0.20 | -0.03 | 0.12 |
| Mediterranean Forests, Woodlands & Scrub | 0.30 | 0.29 | 0.12 | 0 | 0.07 | 0 | 0 | 0.07 | 0.11 |
| Montane Grasslands & Shrublands | 0.35 | 0.26 | 0.80 | 0.06 | 0.73 | 0.86 | 0.64 | 0.43 | 0.44 |
| Temperate Broadleaf & Mixed Forests | 0.22 | 0.22 | 0.19 | 0.001 | 0.01 | 0.09 | 0.18 | 0.03 | 0.09 |
| Temperate Conifer Forests | 0.07 | 0.05 | 0.10 | 0 | -0.02 | 0.22 | -0.08 | 0 | 0.05 |
| Temperate Grasslands, Savannas & Shrublands | 0.18 | 0.14 | 0.23 | 0.03 | 0.11 | 0.01 | 0.19 | 0.09 | 0.05 |
| Tropical & Subtropical Coniferous Forests | 0.28 | 0.32 | 0.25 | 0.61 | 0.43 | 0 | 0.19 | 0.24 | 0.34 |
| Tropical & Subtropical Dry Broadleaf Forests | 0.23 | 0.12 | 0.24 | 0.27 | 0.33 | 0.01 | 0.21 | 0.005 | 0.19 |
| Tropical & Subtropical Grasslands, Savannas & Shrublands | 0.39 | 0.41 | 0.39 | 0.12 | 0.07 | 0.02 | 0.43 | 0.11 | 0.29 |
| Tropical & Subtropical Moist Broadleaf Forests | 0.37 | 0.26 | 0.41 | 0.28 | 0.03 | 0.53 | 0.43 | 0.23 | 0.31 |
| Tundra | 0.06 | 0.06 | 0 | 0 | 0 | 0 | 0 | 0 | 0.05 |

**Figure S1.**
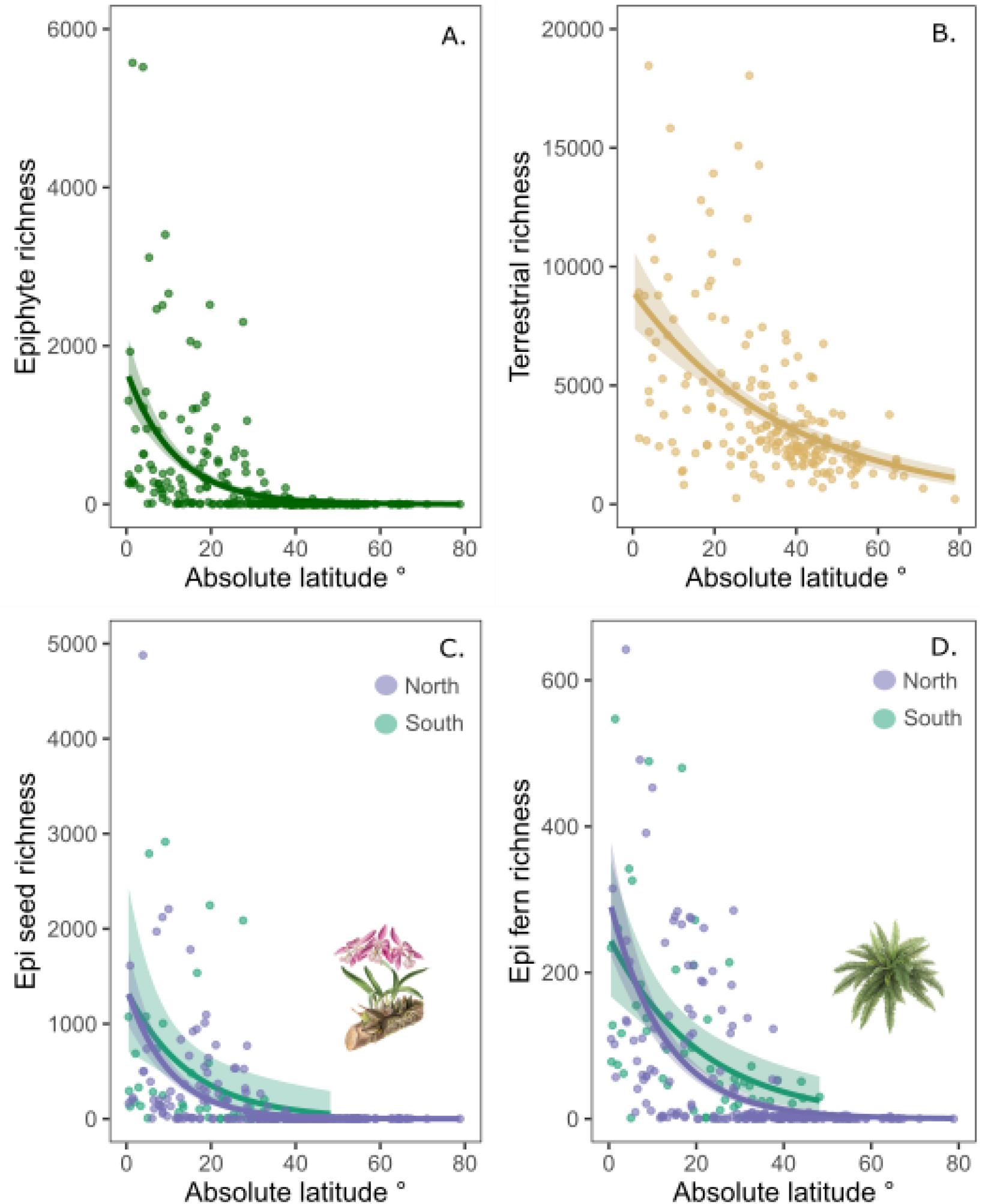
Latitudinal gradients in (A) the total number of epiphyte and (B) terrestrial plant species, and latitudinal asymmetries for (C) seed and (D) pteridophyte epiphyte species between the northern- (purple) and southern hemispheres (blue). Points indicate regions weighted by species richness and lines indicate the strength of the relationship, including 95% confidence intervals.

**Figure S2.**
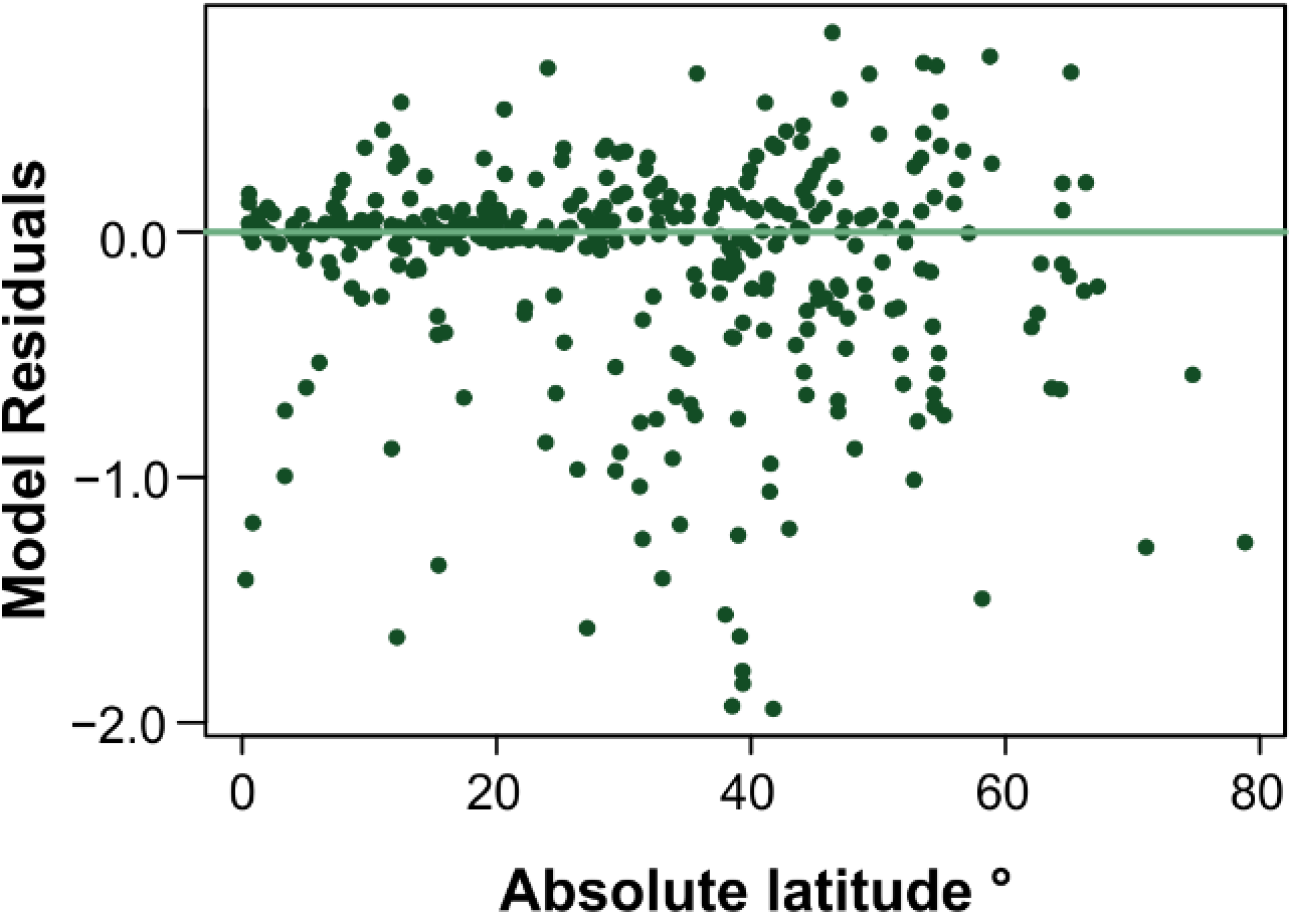
Relationship between residuals of the global epiphyte richness model and absolute latitude.

## Notes

### Competing Interest Statement

The authors have declared no competing interest.

## References

Acebey, A.R., Krömer, T. & Kessler, M. (2017). Species richness and vertical distribution of ferns and lycophytes along an elevational gradient in Los Tuxtlas, Veracruz, Mexico. Flora, 235, 83–91.

Barthlott, W., Hostert, A., Kier, G., Küper, W., Kreft, H., Mutke, J., Rafiqpoor, M.D. & Sommer, J.H. (2007). Geographic patterns of vascular plant diversity at continental to global scales. Erdkunde, 61, 305–315.

Barton, K. (2020). MuMIn: Multi-Model Inference.

Bates, D., Maechler, M., Bolker, B. & Walker, S. (2015). Fitting linear mixed-effects models using lme4. Journal of Statistical Software, 67, 1–48.

Benzing, D.H. (1987). Vascular epiphytism: Taxonomic participation and adaptive diversity. Annals of the Missouri Botanical Garden, 74, 183–204.

Benzing, D.H. (1989). The Evolution of Epiphytism, Vascular Plants as Epiphytes. Springer Berlin, Heidelberg, 76, pp. 15–41.

Benzing, D.H. (1990). Vascular Epiphytes. General Biology and Related Biota. Cambridge University Press, Cambridge.

Benzing, D.H. (1998). Vulnerabilities of tropical forests to climate change: The significance of resident epiphytes. Climate Change, 39, 519–540.

Benzing, D.H. (2004). Vascular Epiphytes, Forest Canopies. Elsevier Academic Press, Burlington, MA, pp. 175–211.

Bivand, R.S. & Wong, D.W.S. (2018). Comparing implementations of global and local indicators of spatial association. TEST, 27, 716–748.

Brummit, R.K. (2001). World Geographical Scheme for Recording Plant Distributions, Edition 2. Biodiversity Information Standards (TDWG) http://www.tdwg.org/standards/109.

Burns, K.C. (2010). How arboreal are epiphytes? A null model for Benzing’s classifications. New Zealand Journal of Botany, 48, 185–191.

Carlucci, M.B., Seger, G.D.S., Sheil, D., Amaral, I.L., Chuyong, G.B., Ferreira, L.V., Galatti, U., Hurtado, J., Kenfack, D., Leal, D.C., Lewis, S.L., Lovett, J.C., Marshall, A.R., Martin, E., Mugerwa, B., Munishi, P., Oliveira, Á.C.A., Razafimahaimodison, J.C., Rovero, F., Sainge, M.N., Thomas, D., Pillar, V.D. & Duarte, L.D.S. (2017). Phylogenetic composition and structure of tree communities shed light on historical processes influencing tropical rainforest diversity. Ecography, 40, 521–530.

Couvreur, T.L.P. (2015). Odd man out: why are there fewer plant species in African rain forests? Plant Systematics and Evolution, 301, 1299–1313.

Danielson, J. & Gesch, D.B. (2011). Global multi-resolution terrain elevation data 2010 (GMTED2010). U.S. Geological Survey Open-File Report 2011-1073, 26 p.

Dawson, J.W. (1986). The vines, epiphytes and parasites of New Zealand forests. Tuatara (Wellington), 28, 43–70.

Dawson, J.W. & Snedon, B. (1969). The New Zealand rain forest: a comparison with tropical rain forest. Pacific Science, 23, 131–147.

Díaz, I.A., Sieving, K.E., Peña-Foxon, M.E., Larraín, J. & Armesto, J.J. (2010). Epiphyte diversity and biomass loads of canopy emergent trees in Chilean temperate rain forests: A neglected functional component. Forest Ecology and Management, 259, 1490–1501.

Dinerstein, E., Olson, D., Joshi, A., Vynne, C., Burgess, N.D., Wikramanayake, E., Hahn, N., Palminteri, S., Hedao, P., Noss, R., Hansen, M., Locke, H., Ellis, E.C., Jones, B., Barber, C.V., Hayes, R., Kormos, C., Martin, V., Crist, E., Sechrest, W., Price, L., Baillie, J.E.M., Weeden, D., Suckling, K., Davis, C., Sizer, N., Moore, R., Thau, D., Birch, T., Potapov, P., Turubanova, S., Tyukavina, A., Souza, N. de, Pintea, L., Brito, J.C., Llewellyn, O.A., Miller, A.G., Patzelt, A., Ghazanfar, S.A., Timberlake, J., Klöser, H., Shennan-Farpón, Y., Kindt, R., Lillesø, J.-P.B., van Breugel, P., Graudal, L., Voge, M., Al-Shammari, K.F. & Saleem, M. (2017). An ecoregion-based approach to protecting half the terrestrial realm. BioScience, 67, 534–545.

Ding, Y., Liu, G., Zang, R., Zhang, J., Lu, X. & Huang, J. (2016). Distribution of vascular epiphytes along a tropical elevational gradient: disentangling abiotic and biotic determinants. Scientific reports, 6, 19706.

Dormann, C., McPherson, J., Araújo, M., Bivand, R., Bolliger, J., Carl, G., Davies, R., Hirzel, A., Jetz, W., Kissling, D., Kühn, I., Ohlemüller, R., Peres-Neto, P., Reineking, B., Schröder, B., Schurr, F. & Wilson, R. (2007). Methods to account for spatial autocorrelation in the analysis of species distributional data: a review. Ecography, 30, 609–628.

Dormann, C.F., Elith, J., Bacher, S., Buchmann, C., Carl, G., Carré, G., Marquéz, J.R.G., Gruber, B., Lafourcade, B. & Leitao, P.J. (2013). Collinearity: a review of methods to deal with it and a simulation study evaluating their performance. Ecography, 36, 27–46.

Ehlers, J., Gibbard, P.L. & Hughes, P.D. (eds.) (2011). Quaternary Glaciations - Extent and Chronology: A Closer Look. Elsevier, Amsterdam.

Gentry, A.H. & Dodson, C.H. (1987). Diversity and biogeography of Neotropical vascular epiphytes. Annals of the Missouri Botanical Garden, 74, 205–233.

Gerstner, K., Dormann, C.F., Václavík, T., Kreft, H. & Seppelt, R. (2014). Accounting for geographical variation in species–area relationships improves the prediction of plant species richness at the global scale. Journal of Biogeography, 41, 261–273.

Givnish, T.J., Barfuss, M.H.J., van Ee, B., Riina, R., Schulte, K., Horres, R., Gonsiska, P.A., Jabaily, R.S., Crayn, D.M., Smith, J.A.C., Winter, K., Brown, G.K., Evans, T.M., Holst, B.K., Luther, H., Till, W., Zizka, G., Berry, P.E. & Sytsma, K.J. (2011). Phylogeny, adaptive radiation, and historical biogeography in Bromeliaceae: insights from an eight-locus plastid phylogeny. American Journal of Botany, 98, 872–895.

Givnish, T.J., Spalink, D., Ames, M., Lyon, S.P., Hunter, S.J., Zuluaga, A., Iles, W.J.D., Clements, M.A., Arroyo, M.T.K., Leebens-Mack, J., Endara, L., Kriebel, R., Neubig, K.M., Whitten, W.M., Williams, N.H. & Cameron, K.M. (2015). Orchid phylogenomics and multiple drivers of their extraordinary diversification. Proceedings of the Royal Society B: Biological Sciences, 28, 20151553.

Göbel, C.Y., Schlumpberger, B.O. & Zotz, G. (2020). What Is a Pseudobulb? Toward a Quantitative Definition. International Journal of Plant Sciences, 181:686–696.

Gotsch, S.G., Nadkarni, N. & Amici, A. (2016). The functional roles of epiphytes and arboreal soils in tropical montane cloud forests. Journal of Tropical Ecology, 32, 455–468.

Hassler, M. (2021). World Ferns. Synonymic checklist and distribution of ferns and lycophytes of the world. https://www.worldplants.de/ferns/.

Henrot, A.-J., François, L., Favre, E., Butzin, M., Ouberdous, M. & Munhoven, G. (2010). Effects of CO 2, continental distribution, topography and vegetation changes on the climate at the Middle Miocene: a model study. Climate of the Past, 6, 675–694.

Hosokawa, T. (1950). Epiphyte-quotient. Bot. Mag. (Tokyo), 63, 18–19.

Karger, D.N., Conrad, O., Böhner, J., Kawohl, T., Kreft, H., Soria-Auza, R.W., Zimmermann, N.E., Linder, H.P. & Kessler, M. (2017). Climatologies at high resolution for the earth’s land surface areas. Scientific Data, 4, 170122.

Karger, D.N., Kessler, M., Lehnert, M. & Jetz, W. (2021) Limited protection and ongoing loss of tropical cloud forest biodiversity and ecosystems worldwide. Nature Ecology & Evolution, 1–9.

Kelly, D.L., O’Donovan, G., Feehan, J., Murphy, S., Drangeid, S.O. & Marcano-Berti, L. (2004). The epiphyte communities of a montane rain forest in the Andes of Venezuela: patterns in the distribution of the flora. Journal of Tropical Ecology, 20, 643–666.

Kier, G., Mutke, J., Dinerstein, E., Ricketts, T.H., Küper, W., Kreft, H. & Barthlott, W. (2005). Global patterns of plant diversity and floristic knowledge. Journal of Biogeography, 32, 1107–1116.

Kissling, W.D., Eiserhardt, W.L., Baker, W.J., Borchsenius, F., Couvreur, T.L.P., Balslev, H. & Svenning, J.-C. (2012). Cenozoic imprints on the phylogenetic structure of palm species assemblages worldwide. Proceedings of the National Academy of Sciences of the United States of America, 109, 7379–7384.

Kreft, H. & Jetz, W. (2007). Global patterns and determinants of vascular plant diversity. Proceedings of the National Academy of Sciences of the United States of America, 104, 5925–5930.

Kreft, H., Jetz, W., Mutke Jens & Barthlott, W. (2010). Contrasting environmental and regional effects on global pteridophyte and seed plant diversity. Ecography, 33, 408–419.

Kreft, H., Köster, N., Küper, W., Nieder, J. & Barthlott, W. (2004). Diversity and biogeography of vascular epiphytes in Western Amazonia, Yasuní, Ecuador. Journal of Biogeography, 31, 1463–1476.

Krömer, T. & Kessler, M. (2006). Filmy ferns (Hymenophyllaceae) as high-canopy epiphytes. Ecotropica, 12, 57–63.

Krömer, T., Kessler, M., Robbert Gradstein, S. & Acebey, A. (2005). Diversity patterns of vascular epiphytes along an elevational gradient in the Andes. Journal of Biogeography, 32, 1799–1809.

Long, J. (2019). jtools: Analysis and Presentation of Scientific Data.

Madison, M. (1977). Vascular epiphytes: their systematic occurrence and salient features. Selbyana, 2, 1–13.

Markgraf, V., McGlone, M. & Hope, G. (1995). Neogene paleoenvironmental and paleoclimatic change in southern temperate ecosystems—a southern perspective. Trends in Ecology & Evolution, 10, 143–147.

McGlone, M.S., Buitenwerf, R. & Richardson, S.J. (2016). The formation of the oceanic temperate forests of New Zealand. New Zealand Journal of Botany, 54, 128–155.

Méndez‐Castro, F.E., Bader, M.Y., Mendieta‐Leiva, G. & Rao, D. (2018). Islands in the trees: A biogeographic exploration of epiphyte‐dwelling spiders. Journal of Biogeography, 45, 2262–2271.

Mendieta-Leiva, G., Porada, P. & Bader, M.Y. (2020). Interactions of epiphytes with precipitation partitioning. Precipitation Partitioning by Vegetation (ed. by I.T. van Stan, E. Gutmann and J. Friesen), pp. 133–146. Springer International Publishing, Cham.

Nadkarni, N.M., Schaefer, D., Matelson, T.J. & Solano, R. (2004). Biomass and nutrient pools of canopy and terrestrial components in a primary and a secondary montane cloud forest, Costa Rica. Forest Ecology and Management, 198, 223–236.

Nervo, M.H., da Silva Coelho, F.V., Windisch, P.G. & Overbeck, G.E. (2016). Fern and lycophyte communities at contrasting altitudes in Brazil’s subtropical Atlantic Rain Forest. Folia Geobotanica, 51, 305–317.

Olson, D.M., Dinerstein, E., Wikramanayake, E.D., Burgess, N.D., Powell, G.V.N., Underwood, E.C., D’amico, J.A., Itoua, I., Strand, H.E., Morrison, J.C., Loucks, C.J., Allnutt, T.F., Ricketts, T.H., Kura, Y., Lamoreux, J.F., Wettengel, W.W., Hedao, P. & Kassem, K.R. (2001). Terrestrial ecoregions of the world: A new map of life on earth. BioScience, 51, 933–938.

Pridgeon, A.M. (1987). The velamen and exodermis of orchid roots. Orchid Biology: Reviews and Perspectives, 4, 141–192.

Proctor, M.C.F. (2007). Ferns, evolution, scale and intellectual impedimenta. New Phytologist, 176, 504–506.

Proctor, M.C.F. (2012). Light and desiccation responses of some Hymenophyllaceae (filmy ferns) from Trinidad, Venezuela and New Zealand: poikilohydry in a light-limited but low evaporation ecological niche. Annals of Botany, 109, 1019–1026.

R Core Team (2020). R: A language and environment for statistical computing, Vienna, Austria.

Ray, N. & Adams, J. (2001). A GIS-based vegetation map of the world at the last glacial maximum (25,000-15,000 BP). Internet Archaeology, 11, 1–44.

Sandel, B., Weigelt, P., Kreft, H., Keppel, G., van der Sande, M.T., Levin, S., Smith, S., Craven, D. & Knight, T.M. (2020). Current climate, isolation and history drive global patterns of tree phylogenetic endemism. Global Ecology and Biogeography, 29, 4–15.

Schimper, A.F.W. (1888). Die Epiphytische Vegetation Amerikas. Jena G. Fischer.

Schimper, A.F.W., Balfour, I.B., Fisher, W.R. & Groom, P. (1903). Plant-geography upon a physiological basis. Clarendon Press, Oxford.

Schuettpelz, E. & Pryer, K.M. (2009). Evidence for a Cenozoic radiation of ferns in an angiosperm-dominated canopy. Proceedings of the National Academy of Sciences of the United States of America, 106, 11200–11205.

Schneider, H., Schuettpelz, E., Pryer, K.M., Cranfill, R., Magallón, S. & Lupia, R. (2004). Ferns diversified in the shadow of angiosperms. Nature, 428, 553–557.

Silvera, K., Santiago, L.S., Cushman, J.C. & Winter, K. (2009). Crassulacean acid metabolism and epiphytism linked to adaptive radiations in the Orchidaceae. Plant Physiology, 149, 1838–1847.

Stuntz, S., Ziegler, C., Simon, U. & Zotz, G. (2002). Diversity and structure of the arthropod fauna within three canopy epiphyte species in central Panama. Journal of Tropical Ecology, 18, 161–176.

Taylor, A., Keppel, G., Weigelt, P., Zotz, G. & Kreft, H. (2021). Functional traits are key to understanding orchid diversity on islands. Ecography, 44, 703–714.

WCSP (2018). World Checklist of Selected Plant Families. Facilitated by the Royal Botanic Gardens, Kew. https://wcsp.science.kew.org/cite.do.

Weigelt, P., König, C. & Kreft, H. (2020). GIFT – A Global Inventory of Floras and Traits for macroecology and biogeography. Journal of Biogeography, 47, 16–43.

WFO (2019). World Flora Online. Published on the Internet; http://www.worldfloraonline.org.

Whittaker, R.J., Triantis, K.A. & Ladle, R.J. (2008). A general dynamic theory of oceanic island biogeography. Journal of Biogeography, 35, 977–994.

Wiens, J.J., Ackerly, D.D., Allen, A.P., Anacker, B.L., Buckley, L.B., Cornell, H.V., Damschen, E.I., Jonathan Davies, T., Grytnes, J.-A., Harrison, S.P., Hawkins, B.A., Holt, R.D., McCain, C.M. & Stephens, P.R. (2010). Niche conservatism as an emerging principle in ecology and conservation biology. Ecology Letters, 13, 1310–1324.

Zotz, G. (2005). Vascular epiphytes in the temperate zones–a review. Plant Ecology, 176, 173–183.

Zotz, G. (2013a). ‘Hemiepiphyte’: a confusing term and its history. Annals of Botany, 111, 1015–1020.

Zotz, G. (2013b). The systematic distribution of vascular epiphytes - a critical update. *Botanical* Journal of the Linnean Society, 171, 453–481.

Zotz, G. (2016). Plants on plants. The biology of vascular epiphytes. Springer, Switzerland.

Zotz, G. & Büche, M. (2000). The epiphytic filmy ferns of a tropical lowland forest - species occurrence and habitat preferences. Ecotropica, 6, 203–206.

Zotz, G., Leja, M., Aguilar-Cruz, Y. & Einzmann, H.J. (2020). How much water is in the tank? An allometric analysis with 205 bromeliad species. Flora, 264, 151557.

Zotz, G., Weigelt, P., Kessler, M., Kreft, H. & Taylor, A. (2021). EpiList 1.0: a global checklist of vascular epiphytes. Ecology, e03326.

